# Automated identification of small drug molecules for Hepatitis C virus through a novel programmatic tool and extensive Molecular Dynamics studies of select drug candidates

**DOI:** 10.1101/2020.07.07.192518

**Authors:** Rafal Madaj, Akhil Sanker, Ben Geoffrey A S, Host Antony David, Shubham Verma, Judith Gracia, Ayodele Ifeoluwa Faleti, Abdulbasit Haliru Yakubu

## Abstract

We report a novel python based programmatic tool that automates the dry lab drug discovery workflow for Hepatitis C virus. Firstly, the python program is written to automate the process of data mining PubChem database to collect data required to perform a machine learning based AutoQSAR algorithm through which drug leads for Hepatitis C virus is generated. The workflow of the machine learning based AutoQSAR involves feature learning and descriptor selection, QSAR modelling, validation and prediction. The drug leads generated by the program are required to satisfy the Lipinski’s drug likeness criteria. 50 of the drug leads generated by the program are fed as programmatic inputs to an In Silico modelling package by the program for fast virtual screening and computer modelling of the interaction of the compounds generated as drug leads and the drug target, a viral Helicase of Hepatitis C. The results are stored automatically in the working folder of the user by the program. The program also generates protein-ligand interaction profiling and stores the visualized images in the working folder of the user. Select protein-ligand complexes associated with structurally diverse ligands having lowest binding energy were selected for extensive molecular dynamics simulation studies and subsequently for molecular mechanics generalized-born surface area (MMGBSA) with pairwise decomposition calculations. The molecular mechanics studies predict In Silico that the compounds generated by the program inhibit the viral helicase of Hepatitis C and prevent the replication of the virus. Thus our programmatic tool ushers in the new age of automatic ease in drug identification for Hepatitis C virus through a programmatic tool that completely automates the dry lab drug discovery workflow. The program is hosted, maintained and supported at the GitHub repository link given below https://github.com/bengeof/Automated-drug-identification-programmatic-tool-for-Hepatitis-C-virus

## Introduction

PubChem is a data repository of chemical compounds, their properties and biological activities [1] which can be programmatically accessed through web API packages such as PUB-REST and python web scrapping techniques [2,3]. Quantitative Structure-Activity Relationship(QSAR) studies are statistical based studies through which drug leads are generated which provide cost cutting advantages in testing and drug discovery for the pharmaceutical industry [4–7]. However the data set required to perform a QSAR study is curated by researchers before performing the statistically study. Our programmatic tool automates this process of data acquisition required to perform a QSAR study to generate drug leads for Hepatitis C virus through programmatic access of PubChem database and python web scrapping techniques [8,9]. The workflow of the QSAR study was also automated through a machine learning based AutoQSAR algorithm. The workflow of a machine learning based AutoQSAR algorithm involves feature learning and descriptor selection, QSAR modelling, validation and prediction [10–12]. The drug leads generated by the program are required to satisfy the Lipinski’s drug likeness criteria as compounds that satisfy Lipinski’s criteria are likely to be an orally active drug in humans [13]. Drug leads generated by the program are fed as programmatic inputs to an In Silico modelling package to computer model the interaction of the compounds generated as drug leads and the drug target of Hepatitis C virus identified with PDB ID : 1A1V. The drug target of Hepatitis C, a viral helicase was identified from the literature with PDB ID : 1A1V [14,15]. The results of the *In Silico* modelling are stored in the working folder of the user. The program also generates protein-ligand interaction profiling and stores the visualized images in the working folder of the user. Select protein-ligand complexes associated with structurally diverse ligands having lowest binding energy obtained from AutoDock-Vina screeening were selected for extensive molecular dynamics simulation studies and subsequently for molecular mechanics generalized-born surface area (MMGBSA) with pairwise decomposition calculations. The molecular mechanics studies predict that the compounds generated by the program inhibit the viral helicase of Hepatitis C and prevent the replication of the virus. Therefore our new programmatic tool ushers in a new age of automatic ease in drug identification for Hepatitis C virus through a fully automated QSAR and an automated In Silico modelling of the drug leads generated by the autoQSAR algorithm.

Our work is distinguished from previous attempts of virtual screening of large ligand libraries in way that we employ programmatic techniques as compared to other works that do not [16]. However as compared to recent data drive machine learning based approaches to drug discovery [17,18] that use pre-downloaded data sets we deploy a real time data mining which makes a case for a dynamic approach to drug lead generation and the results of the program are reflective of PubChem data library at the instant the program is run and thus approaches drug discovery from a dynamic approach in an age Big Data and constantly growing data libraries. We also add to the existing richness of the novelty of methods [18–20] in data driven drug discovery in the following way. Our programmatic tool couples the drug leads generated by the AutoQSAR algorithm as programmatic inputs to an *In Silico* modelling package and programmatically profiles the protein-ligand interaction and stores the results in the working folder of the user. While adding to the existing richness in data driven machine learning based drug discovery methods, our work also adds new scientific findings to existing literature as the drug target of Hepatitis C virus chosen for the study has not been approached by data driven machine learning based methods.

## Methods and Techniques

The workflow of the programmatic tool implementing programmatic data mining, AutoQSAR and automated In Silico modelling for identification of drugs against Hepatitis C virus is shown in Fig.1. The first process involved in programmatic workflow is the data mining of PubChem database to automate the process of data acquisition to implement a machine learning based AutoQSAR algorithm that automates the process of drug lead generation for Hepatitis C virus. The programmatic access to PubChem is accomplished through python commands [8, 9]. The next process in the workflow involves implementing the machine learning based AutoQSAR algorithm for drug lead generation. The drug leads are generated by the AutoQSAR algorithm through the workflow that involves feature learning, descriptor selection, QSAR modelling, validation and prediction [10–12]. Based on the validated QSAR model PubChem compound library is screened for drug lead generation based on the validated QSAR model. The drug leads generated by the program are filtered through the Lipinski’s criteria [13].

**Fig. 1.**
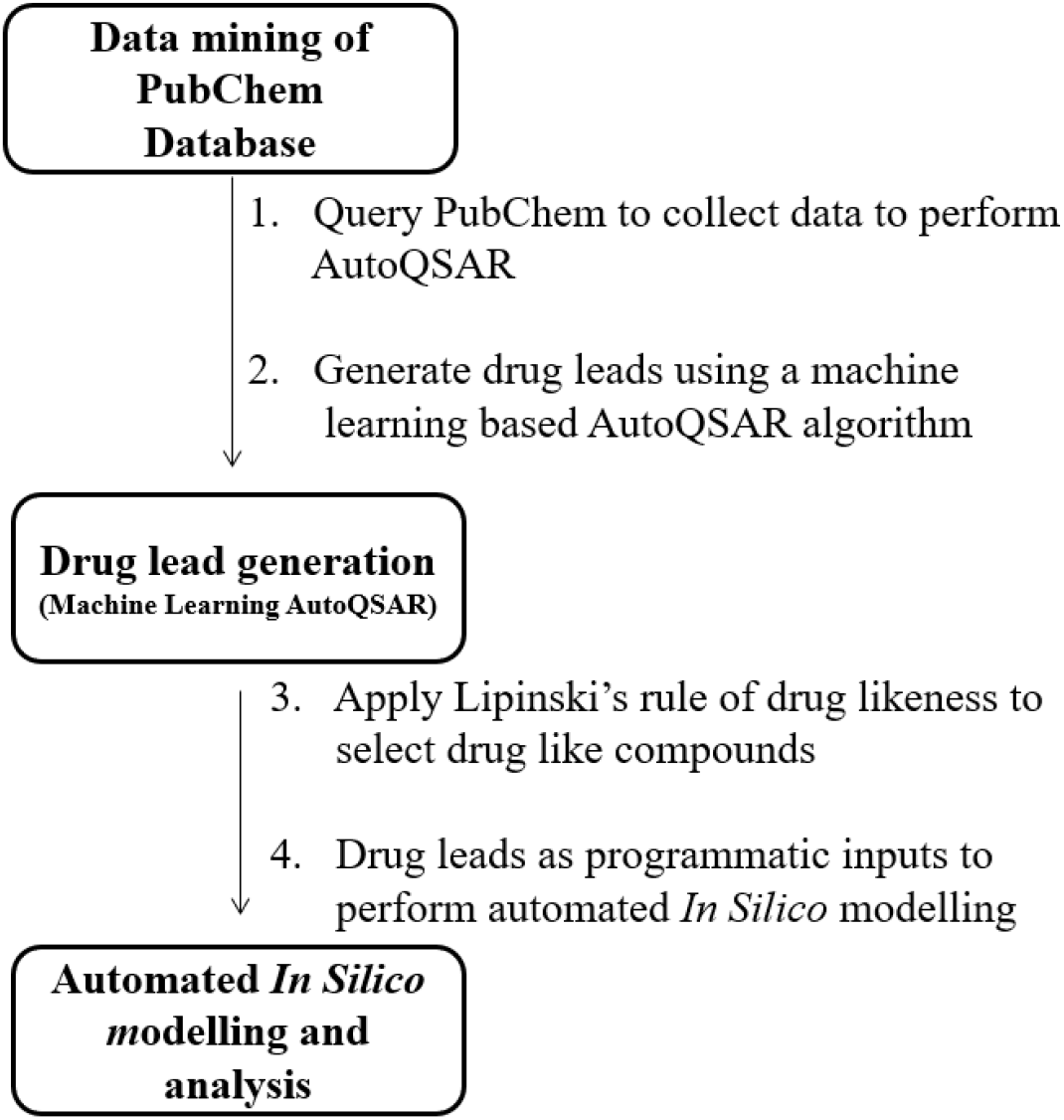
Workflow of the programmatic tool to automate the drug discovery for Hepatitis C virus

Running the program requires no more programming knowledge than running the python executable file in python 3 environment in Linux OS along with some python dependency packages installed such as:

pandas

biopandas

numpy

matplotlib

scikit-learn

seaborn

selenium (along with selenium’s driver for firefox browser)

Other additional dependencies for automated *In Silico* modelling

openbabel 2.4.1

mgltools 1.5.4

autodock-vina 1.1.2-4

The program is hosted, maintained and supported at the GitHub repository link given below https://github.com/bengeof/Automated-drug-identification-programmatic-tool-for-Hepatitis-C-virus

The running of the program requires a stable internet connection and the run time of the program is expected to be a few hours however it is expected to vary based on CPU and internet speed. The program prints out the PubChem CIDs of compounds identified as drug leads for Hepatitis C (shown in Fig.2).

**Fig.2.**
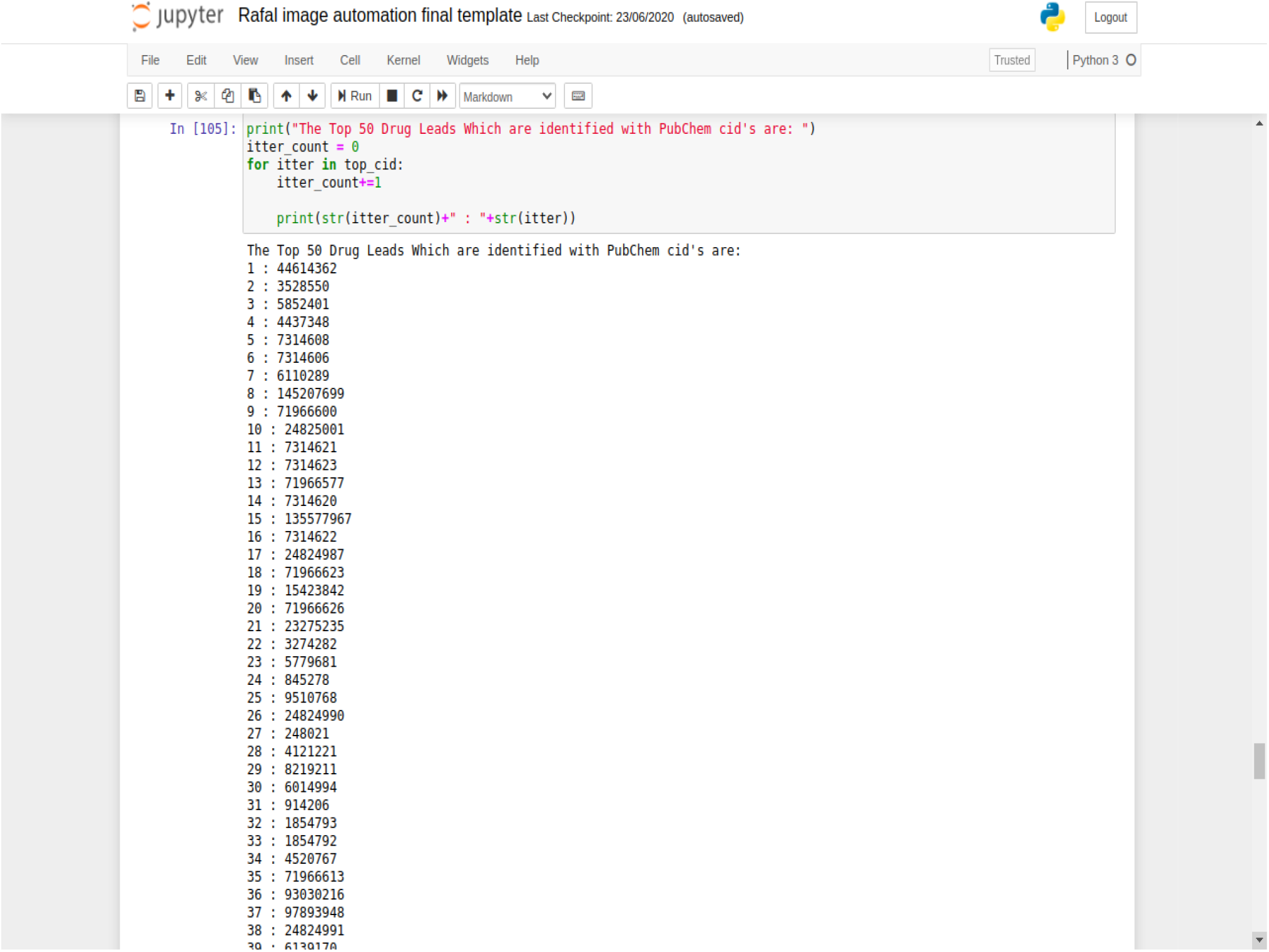
PubChem CIDs of compounds identified as drug leads against Hepatitis C virus

The crystal structure of the Hepatitis C viral helicase identified as the drug target was obtained from the RCSB-PDB database [21] with PDB ID : 1A1V. The ligand and the protein files were prepared for AutoDock process using AutoDockTools(ADT) scripts and the protein drug target files are to be kept in the working folder of the user and can be downloaded from the GitHub repository. The structure of the drug lead compounds generated by the program were programmatically downloaded from PubChem and programmatically prepared for molecular docking using AutoDockTools ligand preparation scripts. The virtual screening using AutoDock Vina [22,23] was initiated programmatically through the program and the interaction between the drug targets proteins and lead drug compounds also automatically profiled and the visualized image of the protein-ligand interaction is saved in the working folder of the user by the program [24,25]. The program also allows the user to control the number of compounds that must proceed for *In Silico* analysis in the following way as shown in Fig.3 and thus allowing the user to control the program as required by the research needs. The program stores the results of the *In Silico* modelling in the working folder of the user and accesses them programmatically as shown in Fig.4 to produce proteinligand interaction profiling images and stores the results in the working folder of the user.

**Fig.3.**
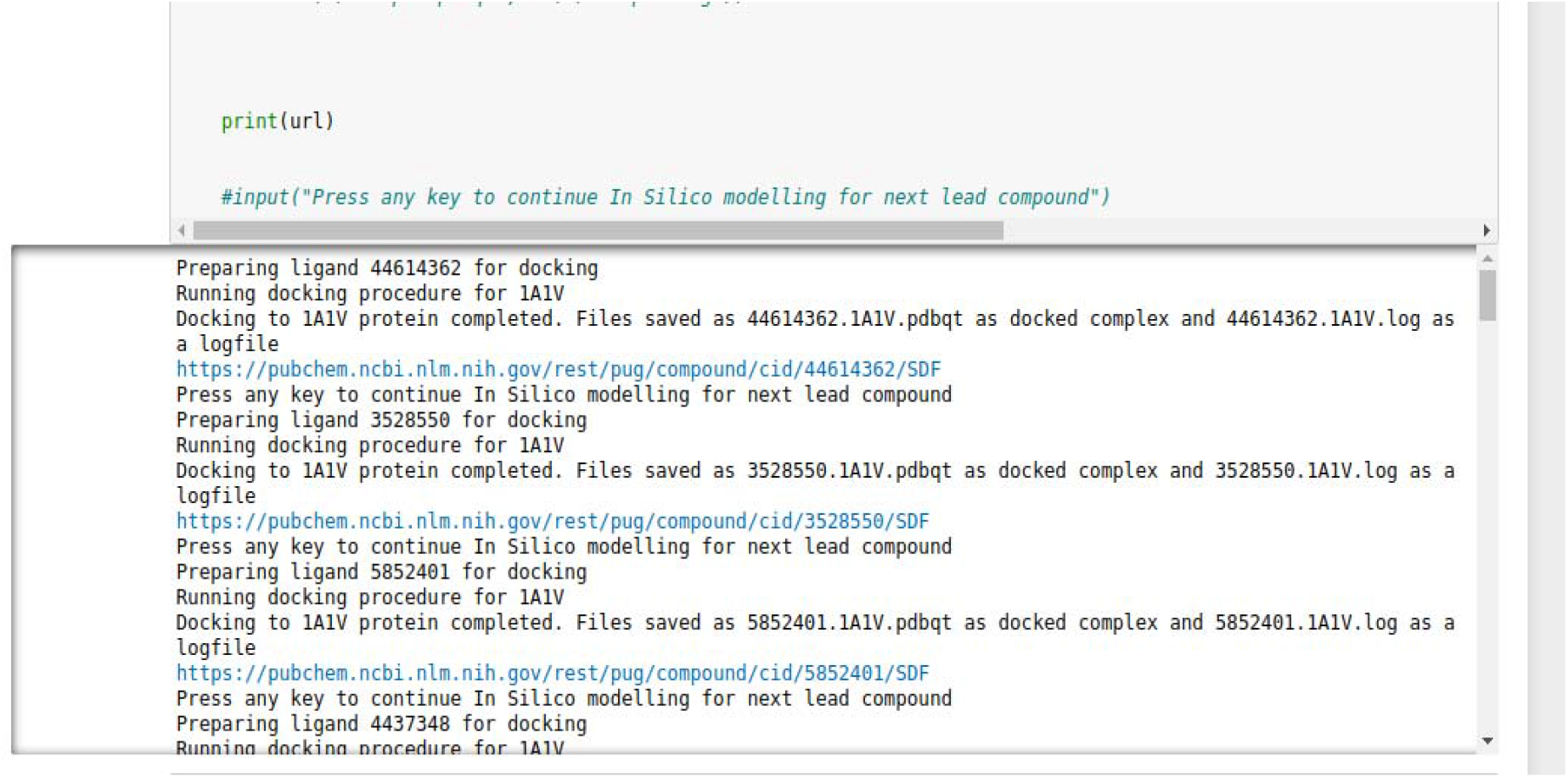
Automated *In Silico* modelling

**Fig. 4.**
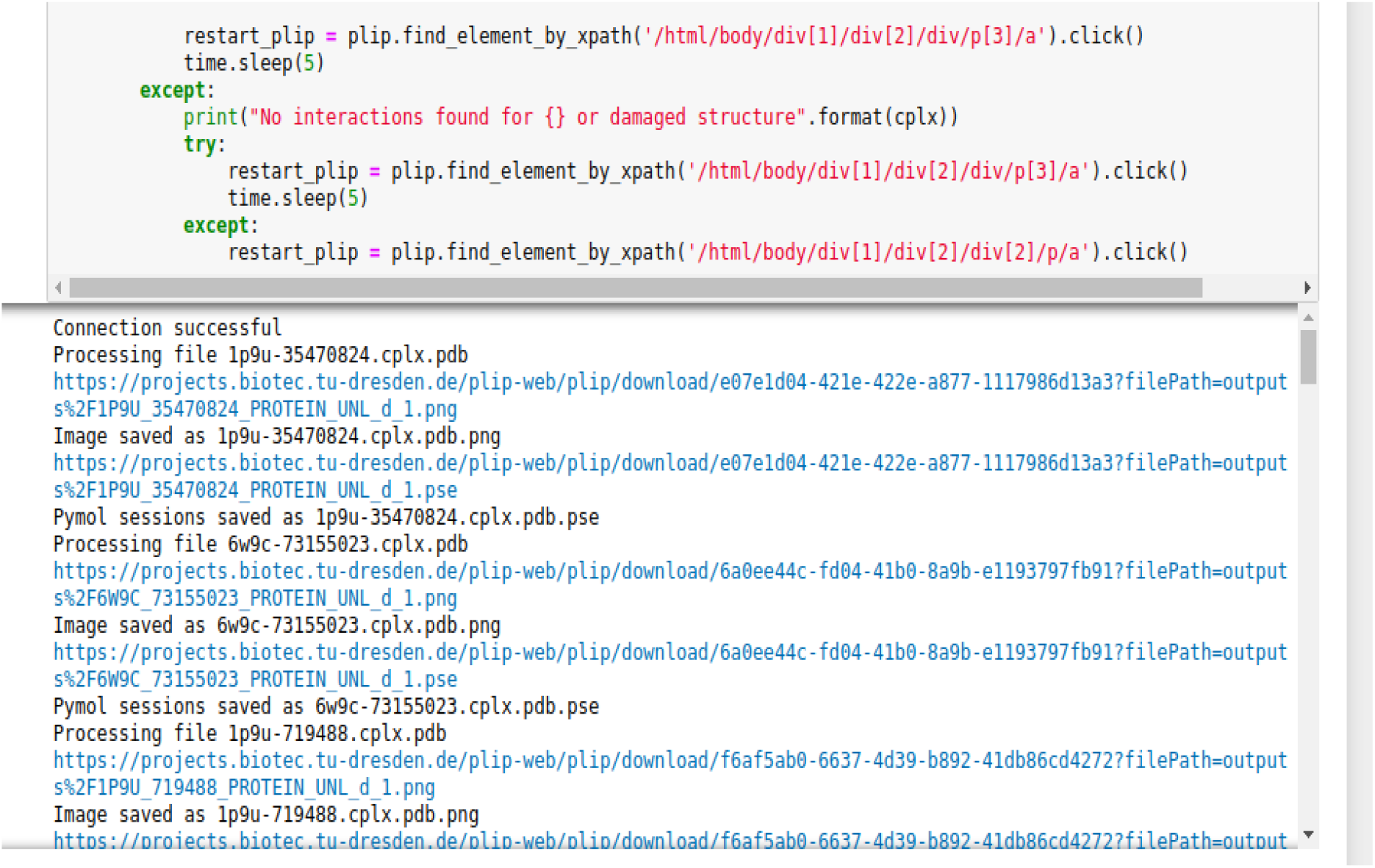
Automated Protein-Ligand profiling of interaction between lead compounds and Hepatitis C drug target protein

Select protein-ligand complexes associated with structurally diverse ligands having lowest binding energy obtained from AutoDock-Vina screening were selected for extensive molecular dynamics simulation studies and subsequently for molecular mechanics generalized-born surface area (MMGBSA) with pairwise decomposition calculations. The ligands were cut out from the complexes and optimized using AmberTools18 from Amber18 suite, followed by partial charges calculation according to AM1-BCC level of theory. The topology and input coordinates were created using tleap. The protein was described using ff14SB force field, ligand using GAFF and water molecules by TIP3P. The system was placed inside truncated octahedral box with 16 A boundary, solvated and neutralized. Minimization encompassed 20000 steps, 10000 steepest descent algorithm and the rest using conjugate gradients. The heating encompassed 50 to 300 K increase in the temperature and lasted 100 ps. Restraints on protein backbone of 4 kcal/mol*A^o^ were applied. The production run were performed under NPT ensemble for 20 ns, out of which 10 ns were truncated for equilibration purposes. The simulation runs were put under periodic boundary conditions, non-bonded interactions were evaluated with the Particle Mesh Ewald method with cut-off of 9 A^o^. Langevin thermostat and Monte Carlo Barostat were used for temperature and pressure maintenance. All described simulations were done using pmemd module of AMBER 18, utilizing CPU for minimization and GPU for heating, equilibration and production. For each ligand, the simulations were repeated ten times. Subsequently, the trajectories were merged and clustered. The clusters encompassing ligand within the binding cavity were selected for MMGBSA calculations and then pairwise decomposition. The calculation were performed on the Intel^®^ CoreTM i9-9900KF CPU @ 3.60GHz × 16 with 32GB @ 2666MHz with GeForce RTX 2070 SUPER/PCIe/SSE2 on the Ubuntu 20.04 Focal Fossa.

## Results and Discussion

The python program was run in Python 3 environment with the dependency packages mentioned in the methodology section. The program prints out the PubChem CIDs of the top 50 compounds identified as Hepatitis C virus drug leads by the program and is shown in Fig.2. This is done by the program through automated programmatic data mining of PubChem database to collect data required to perform a machine learning based AutoQSAR algorithm on the dataset to generate the drug leads for Hepatitis C virus. The drug leads generated by the program for the Hepatitis C virus are useful to screen PubChem database which is over a billion compounds and the generated drug leads are useful to further pursue *In Silico, In Vitro and In Vivo* testing and is expected to save computational and experimental testing costs for the pharmaceutical industry. The drug leads generated by the program were required to satisfy the Lipinski’s criteria of drug likeness as compounds that satisfy the Lipinski’s criteria are likely to be orally active drug in humans. The structure of the compounds identified as drug leads were programmatically downloaded from PubChem by the program and they were fed as programmatic ligand input files after ligand preparation via ADT scripts to a *In Silico* modelling package used widely known as AutoDock-Vina. Therefore the study of the interaction of the drug lead compounds and the viral helicase of Hepatitis C virus identified as the drug target was automated through the programmatic inputs given to AutoDock-Vina in the program. The results of the virtual screening for the top 50 drug lead compounds are given in Table 1. The protein-ligand interaction of select protein-ligand complexes associated with structurally diverse ligands having lowest binding energy are shown in Fig. 5a,b,c&d. Select protein-ligand complexes associated with structurally diverse ligands having lowest binding energy obtained from AutoDock-Vina screening were selected for extensive molecular dynamics simulation studies and subsequently for molecular mechanics generalized-born surface area (MMGBSA) with pairwise decomposition calculations. The results of the molecular dynamics studies are shown in Table 2 and they predict that the compounds generated by the program inhibit the viral helicase of Hepatitis C *In Silico* and prevent the replication of the virus.

**Fig.5a.**
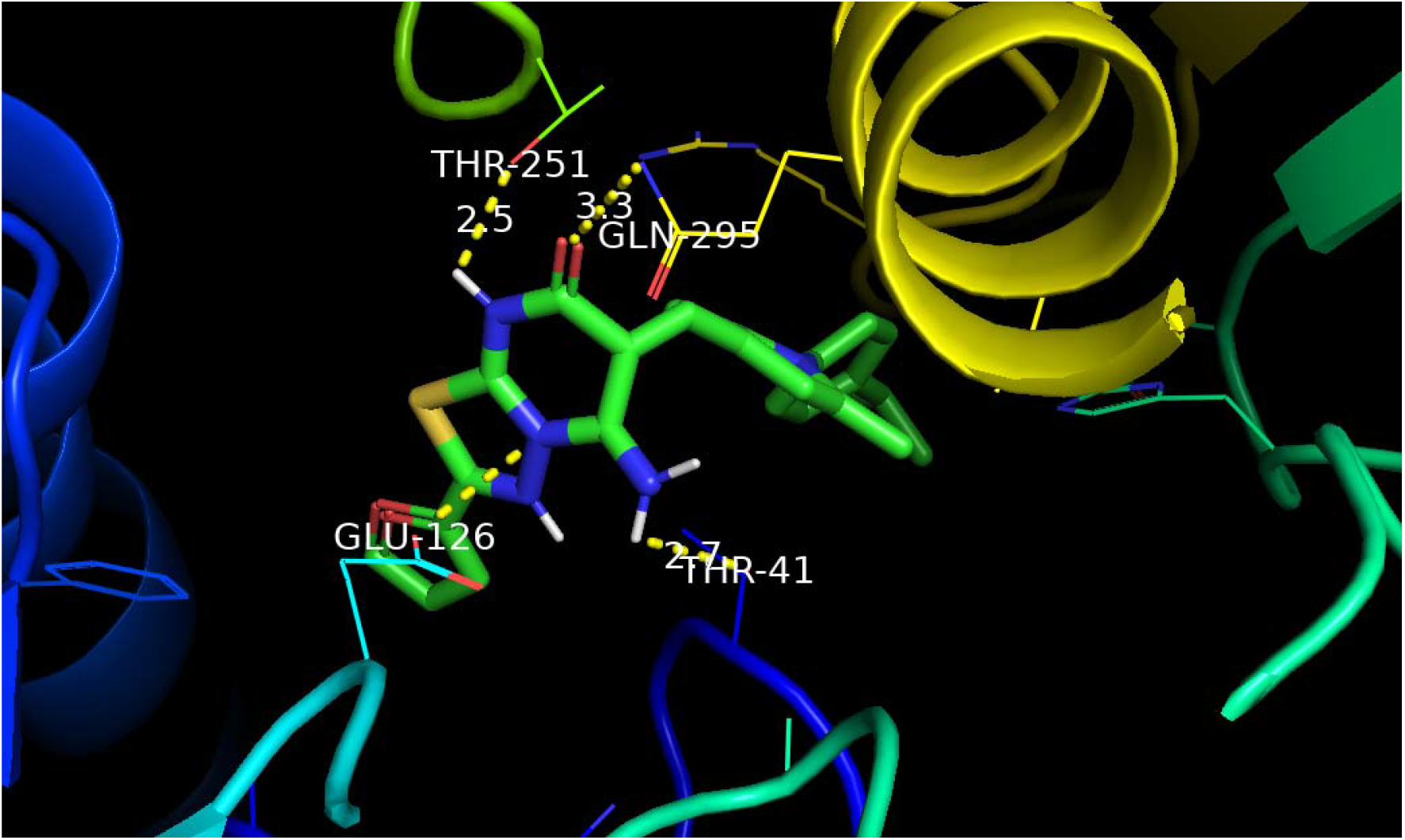
Interaction of drug target of Hepatitis C and compound identified with PubChem CID 362

**Fig. 5b.**
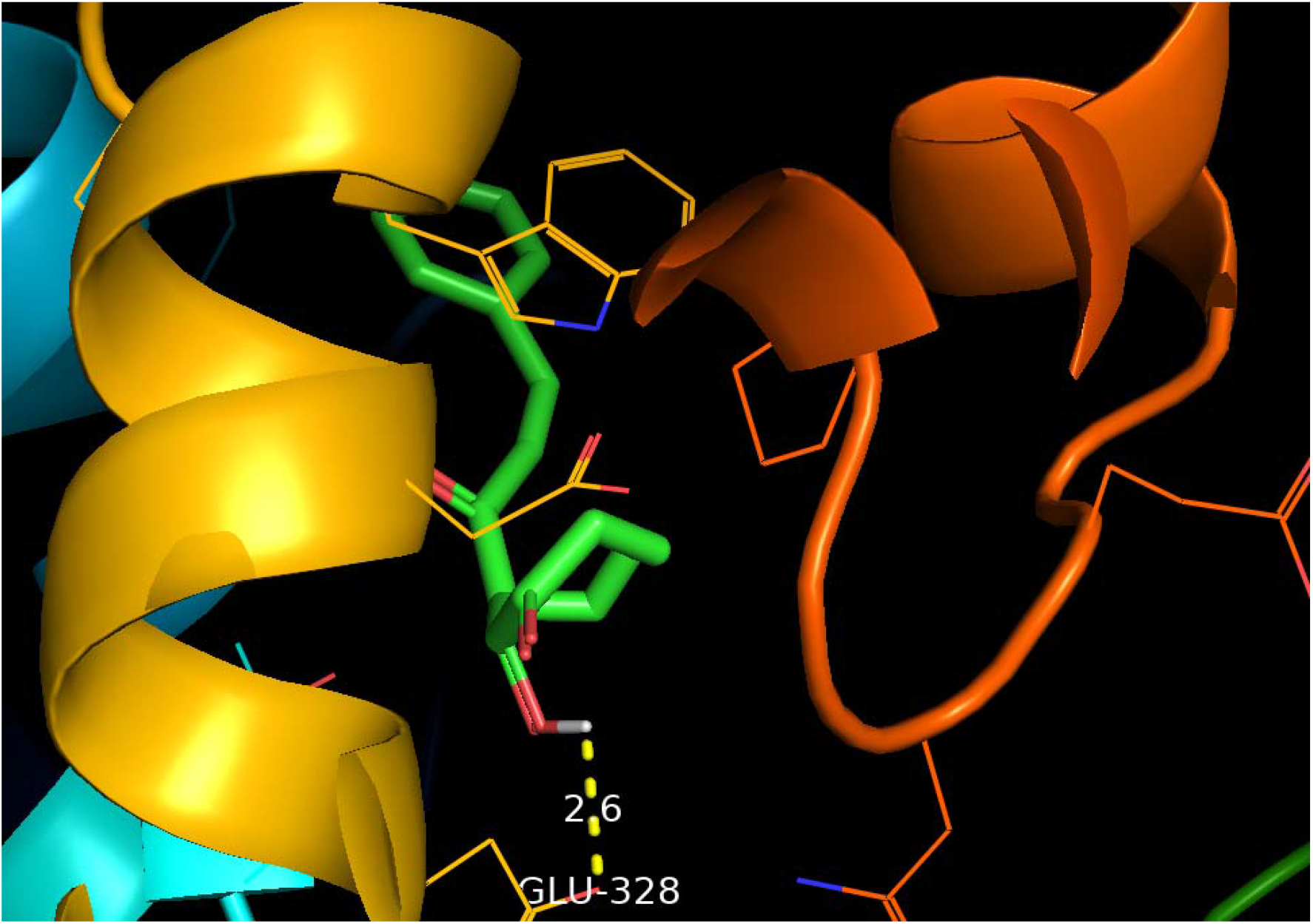
Interaction of drug target of Hepatitis C and compound identified with PubChem CID 842

**Fig. 5c.**
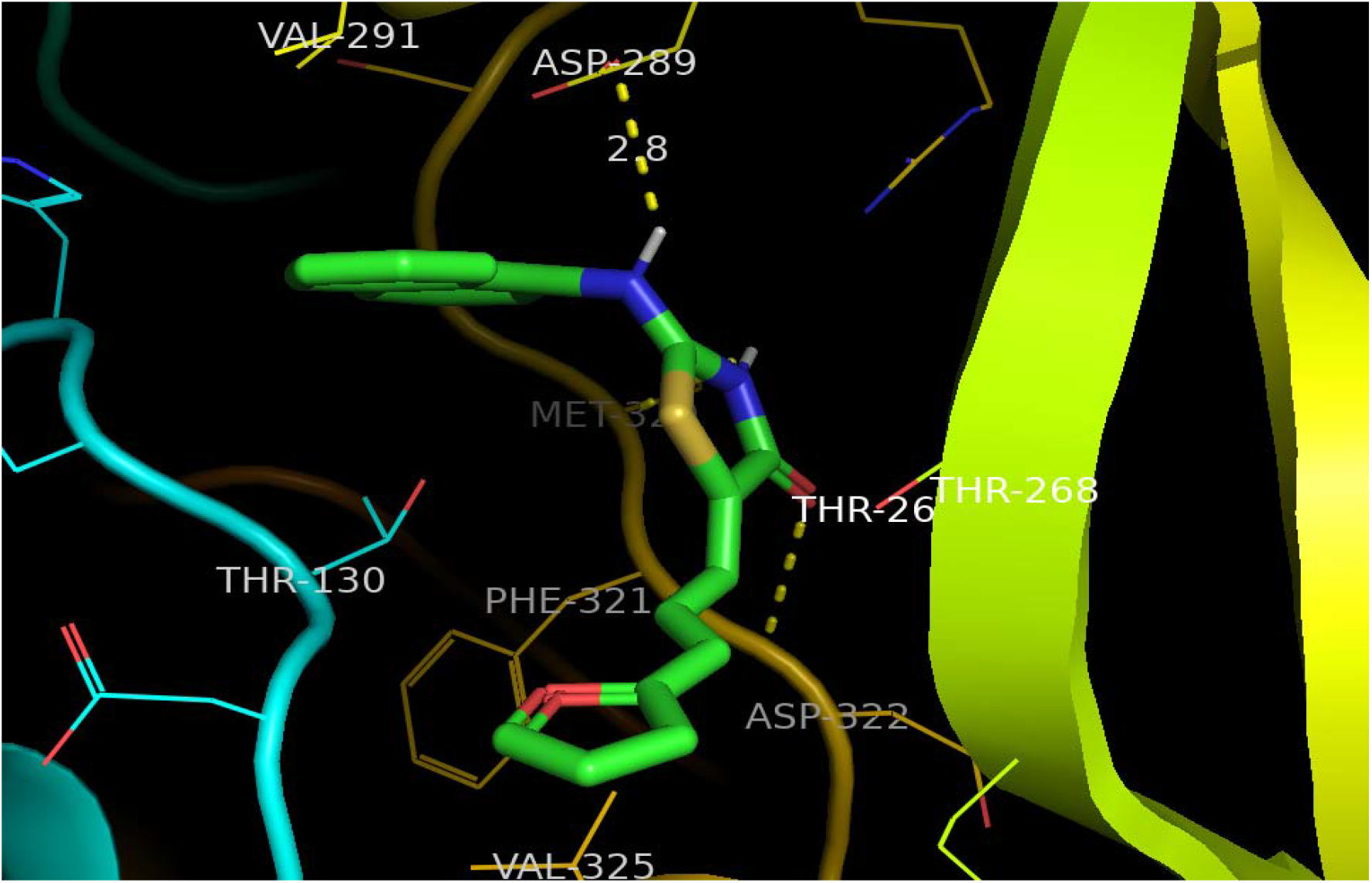
Interaction of drug target of Hepatitis C and compound identified with PubChem CID 602

**Fig. 5d.**
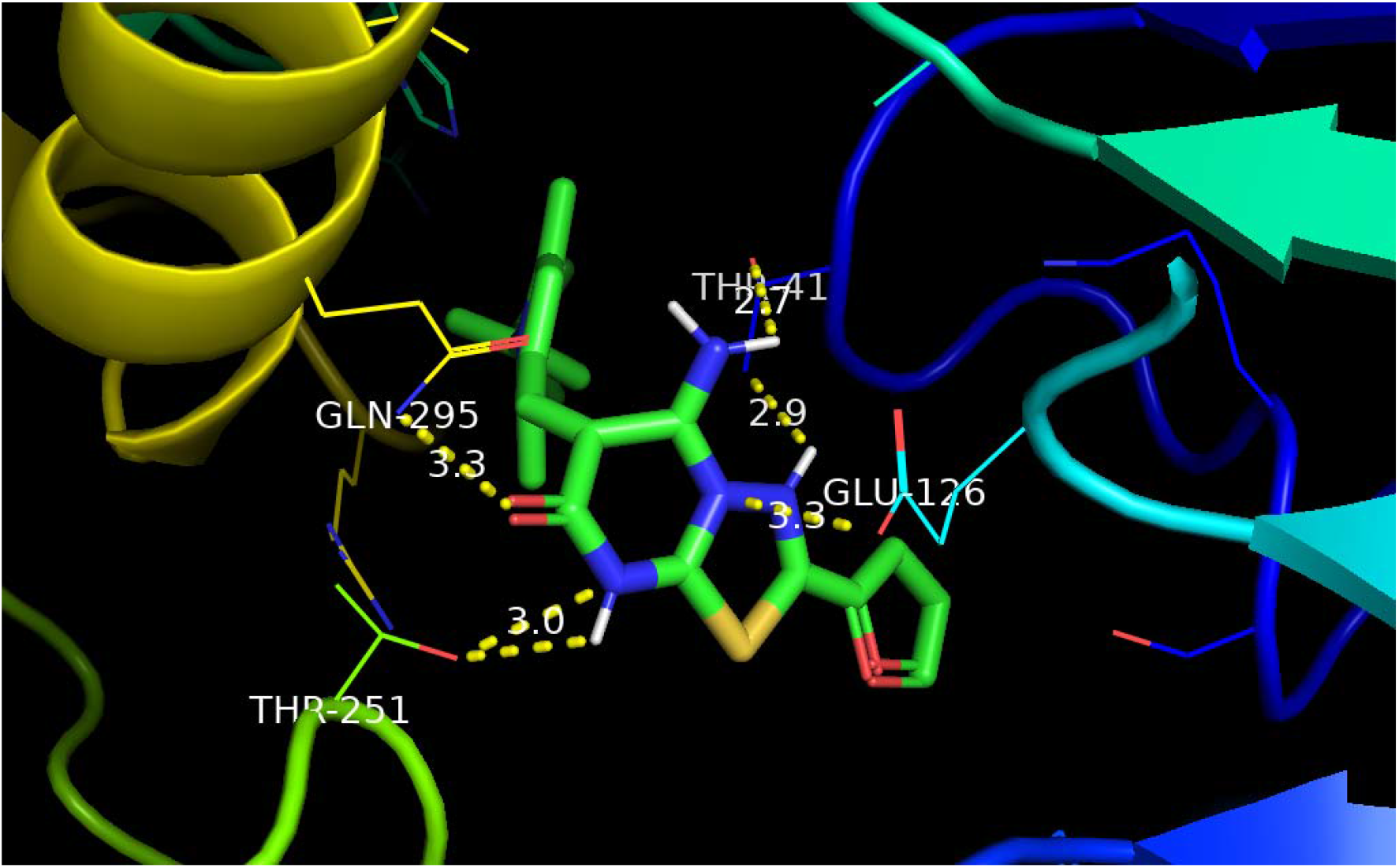
Interaction of drug target of Hepatitis C and compound identified with PubChem CID 792

**Table 1.**
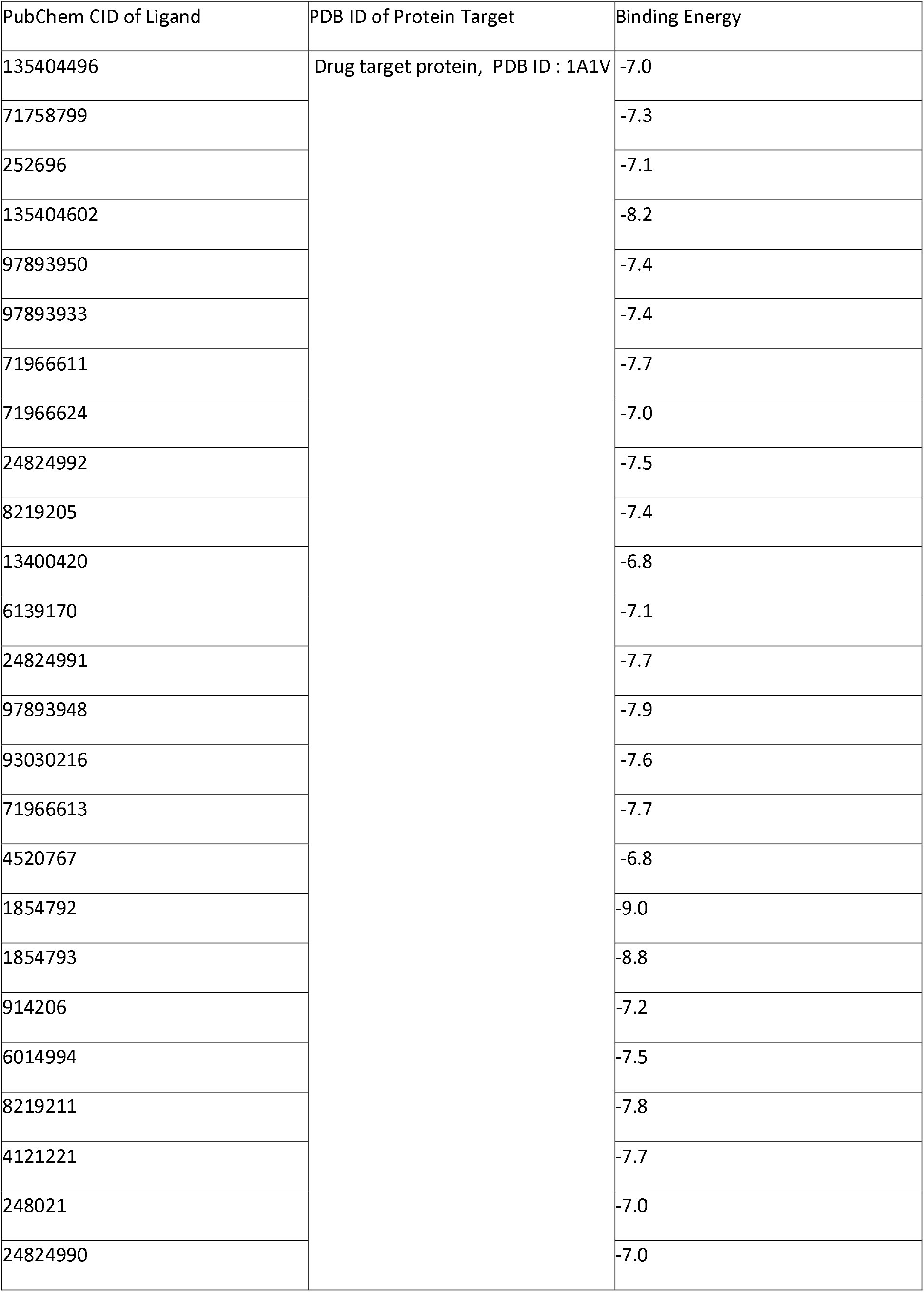
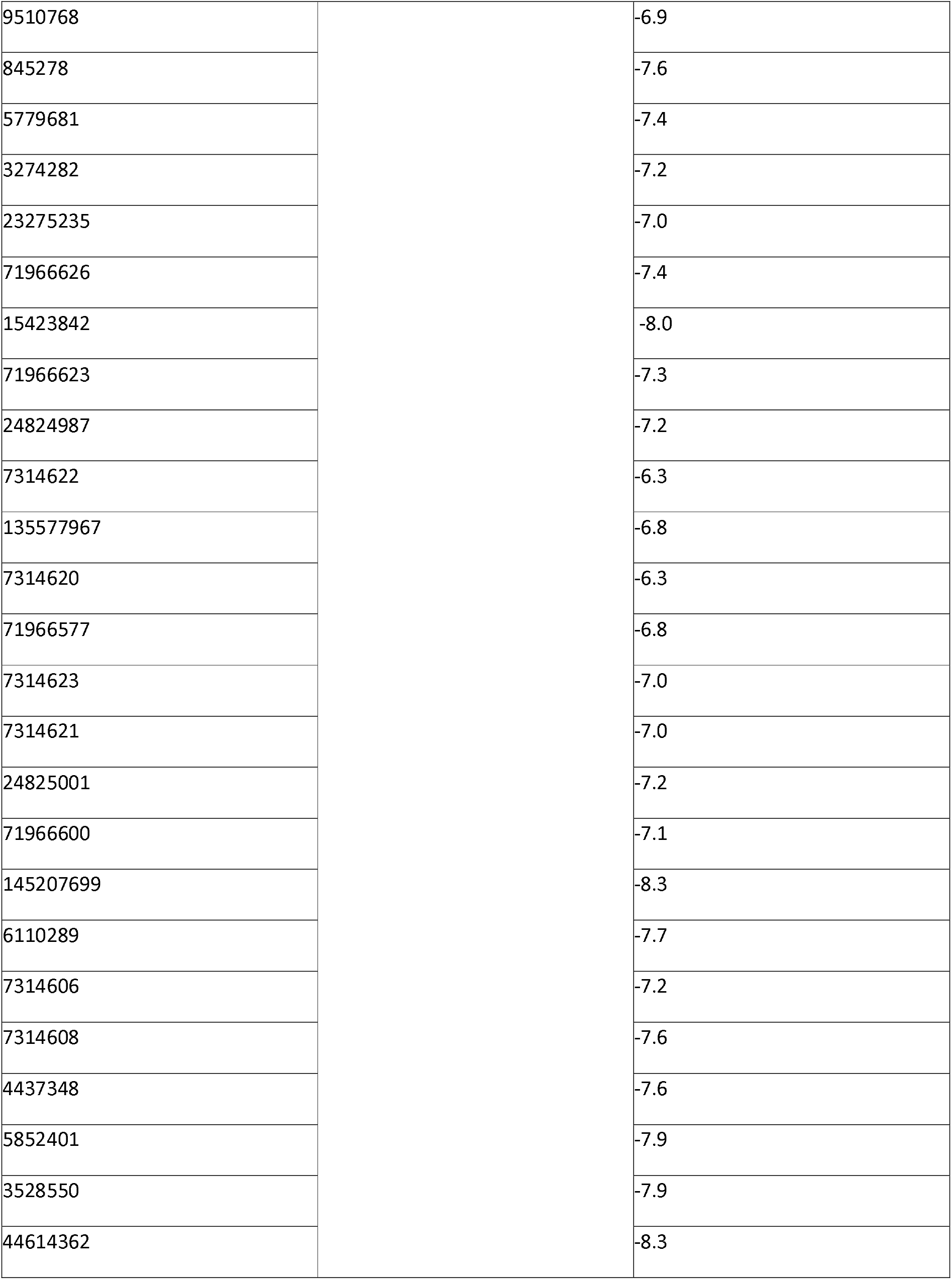
Interaction of lead drug compounds and Hepatitis C virus drug target protein

| PubChem CID of Ligand | PDB ID of Protein Target | Binding Energy |
| --- | --- | --- |
| 135404496 | Drug target protein, PDB ID : 1A1V | -7.0 |
| 71758799 |  | -7.3 |
| 252696 |  | -7.1 |
| 135404602 |  | -8.2 |
| 97893950 |  | -7.4 |
| 97893933 |  | -7.4 |
| 71966611 |  | -7.7 |
| 71966624 |  | -7.0 |
| 24824992 |  | -7.5 |
| 8219205 |  | -7.4 |
| 13400420 |  | -6.8 |
| 6139170 |  | -7.1 |
| 24824991 |  | -7.7 |
| 97893948 |  | -7.9 |
| 93030216 |  | -7.6 |
| 71966613 |  | -7.7 |
| 4520767 |  | -6.8 |
| 1854792 |  | -9.0 |
| 1854793 |  | -8.8 |
| 914206 |  | -7.2 |
| 6014994 |  | -7.5 |
| 8219211 |  | -7.8 |
| 4121221 |  | -7.7 |
| 248021 |  | -7.0 |
| 24824990 |  | -7.0 |
| 9510768 |  | -6.9 |
| 845278 |  | -7.6 |
| 5779681 |  | -7.4 |
| 3274282 |  | -7.2 |
| 23275235 |  | -7.0 |
| 71966626 |  | -7.4 |
| 15423842 |  | -8.0 |
| 71966623 |  | -7.3 |
| 24824987 |  | -7.2 |
| 7314622 |  | -6.3 |
| 135577967 |  | -6.8 |
| 7314620 |  | -6.3 |
| 71966577 |  | -6.8 |
| 7314623 |  | -7.0 |
| 7314621 |  | -7.0 |
| 24825001 |  | -7.2 |
| 71966600 |  | -7.1 |
| 145207699 |  | -8.3 |
| 6110289 |  | -7.7 |
| 7314606 |  | -7.2 |
| 7314608 |  | -7.6 |
| 4437348 |  | -7.6 |
| 5852401 |  | -7.9 |
| 3528550 |  | -7.9 |
| 44614362 |  | -8.3 |

**Table 2.** Molecular Dynamics studies

| Method | Molecular docking | Molecular dynamics |  |  |
| --- | --- | --- | --- | --- |
| Feature | Lowest free energy of binding $\Delta G$ [kcal/mol] | % of time spent inside binding cavity | Total relative binding free energy $\Delta\Delta G$ [kcal/mol] | Most important resids (more than -1 kcal/mol contribution) |
| 1854792 | -9.0 | 69.00% | $-26.4 \pm 5.8$ | PRO40, THR41, GLU126, ALA158, HID168, ASN170, THR251, GLN295, GLY298, ARG 299, ARG 302 |
| 15423842 | -8.0 | 86.70% | $-14.8 \pm 5.8$ | THR98, GLY90, THR104, GLY106, LYS107, ALA110, ARG228, TRP336, TYR337 |
| 135404602 | -8.2 | 89.00% | $-24 \pm 4.6$ | THR130, CYS266, THR268, GLN269, LEU 286, GLN288, SER292, ARG316, PHE321, ASP322, SER324, VAL 325, GLU328 |
| 44614362 | -8.3 | 91.10% | $-14 \pm 3.0$ | LYS107, ALA110, TRP336, TYR337, LYS386 |

Therefore usefulness of the tool is demonstrated in reducing the complexity of virtual screening for drugs against Hepatitis C virus achieved through programmatic automation of data mining of PubChem to collect data required to implement a machine learning based AutoQSAR algorithm for automatic drug lead generation. The program requires that the drug leads generated by the program were required to satisfy the Lipinski’s rule drug-likeness criteria. Further, the program automates the In Silico modelling of the interaction of the compounds generated as drug leads and the drug target of Hepatitis C virus and stores the results in the working folder of the user. Thus, the program helps achieve complete automation in identifying drugs against Hepatitis C virus which further has to be examined for drug potential through experimental testing such as *In Vitro* and *In Vivo* testing.

## Conclusion and future scope

Thus the presented work is an attempt to automate the dry lab drug discovery workflow for discovering drugs for Hepatitis C virus by a python program that automates the process of data mining PubChem database to collect data required to perform a machine learning based AutoQSAR algorithm through which drug leads for Hepatitis C virus are generated. The data acquisition from PubChem was carried out through python web scrapping techniques. The workflow of a machine learning based AutoQSAR involves feature learning and descriptor selection, QSAR modelling, validation and prediction. The drug leads generated by the program are required to satisfy the Lipinski’s drug likeness criteria. Drug leads generated by the program are fed as programmatic inputs to an In Silico modelling package to computer model the interaction of the compounds generated as drug leads and the drug target, a viral helicase (PDB ID :1A1V). The results are stored in the working folder of the user. The program also generates protein-ligand interaction profiling and stores the visualized images in the working folder of the user. Select protein-ligand complexes associated with structurally diverse ligands having lowest binding energy obtained from AutoDock-Vina screeening were selected for extensive molecular dynamics simulation studies and subsequently for molecular mechanics generalized-born surface area (MMGBSA) with pairwise decomposition calculations. The molecular mechanics studies predict that the compounds generated by the program inhibit the viral helicase of Hepatitis C and prevent the replication of the virus. Therefore our new programmatic tool ushers in a new age of automatic ease in drug identification for Hepatitis C virus through a fully automated QSAR and an automated In Silico modelling of the drug leads generated by the AutoQSAR algorithm. While the compounds identified through the automated workflow must be test experimentally by the experimental research community for their drug potential against Hepatitis C virus, there is still a lot of scope to make the automation algorithm more self-aware of technical nuances which will help increase its accuracy in drug identification which we bring to the attention of the computational research community for their scholarly attention and efforts on the same.

The authors declare we have no conflict of Interest to disclose and that the work is original and is not being considered for publication elsewhere

